# Replication and Refinement of Brain Age Model for adolescent development

**DOI:** 10.1101/2023.08.16.553472

**Authors:** Bhaskar Ray, Jiayu Chen, Zening Fu, Pranav Suresh, Bishal Thapaliya, Britny Farahdel, Vince D. Calhoun, Jingyu Liu

## Abstract

The discrepancy between chronological age and estimated brain age, known as the brain age gap, may serve as a biomarker to reveal brain development and neuropsychiatric problems. This has motivated many studies focusing on the accurate estimation of brain age using different features and models, of which the generalizability is yet to be tested. Our recent study has demonstrated that conventional machine learning models can achieve high accuracy on brain age prediction during development using only a small set of selected features from multimodal brain imaging data. In the current study, we tested the replicability of various brain age models on the Adolescent Brain Cognitive Development (ABCD) cohort. We proposed a new refined model to improve the robustness of brain age prediction. The direct replication test for existing brain age models derived from the age range of 8-22 years onto the ABCD participants at baseline (9 to 10 years old) and year-two follow-up (11 to 12 years old) indicate that pre-trained models could capture the overall mean age failed precisely estimating brain age variation within a narrow range. The refined model, which combined broad prediction of the pre-trained model and granular information with the narrow age range, achieved the best performance with a mean absolute error of 0.49 and 0.48 years on the baseline and year-two data, respectively. The brain age gap yielded by the refined model showed significant associations with the participants’ information processing speed and verbal comprehension ability on baseline data.

## 1 Introduction

As we grow old, our brain develops and changes parallelly throughout our life. Extracting insightful information from neuroimaging data underlying brain development has become a widely studied research area recently [1–3]. An individual’s brain age is referred to as biological age, which cannot be measured directly but can be modeled through neuroimaging data. Brain age models are usually trained with brain imaging features of magnetic resonance imaging (MRI) or other types of neuroimaging resources from a healthy or typical development population. The brain age gap is computed by the difference between chronological age and estimated brain age. Recently brain age gap has been utilized as a potential biological indicator for identifying several neurodegenerative disorders such as Alzheimer’s and Parkinson’s at their early stage [4–6], or for understanding brain development trajectory or abnormal aging process [7–9].

Multiple studies have been conducted using various methods such as voxel-based, region-based, and surface-based techniques to obtain features from different modalities to accurately predict brain age. The early studies of brain age were mainly focused on using traditional machine learning techniques such as regression-based approaches to estimate brain age [10–13]. For instance, the authors of [11] applied a support vector regression (SVR) model with 1,116,006 features derived from multimodal structural imaging maps of 621 subjects from the Philadelphia Neurodevelopmental Cohort (PNC) cohort aged 8-22 years and achieved a mean absolute error (MAE) of 1.22 years on estimated brain age. Lund et al. [13] used a single modality, resting-state fMRI imaging data, to analyze the brain development of adolescents from the PNC cohort and measured the relationship between individuals’ cognitive ability and brain age gap. Based on the shrinkage estimation of the regression model, the MAE was 2.43 years with a correlation of 0.60 between the actual age and the predicted age. When the same model was tested on an independent sample of subjects in the same age group (8-22 years) from the healthy brain network cohort, it achieved an MAE of 3.46 years and a correlation value of r = 0.58.

Deep learning models have recently gained popularity in the brain age research field due to their ability to produce consistent promising results in various imaging modalities [14–19]. Peng et al. [14] developed a simple, fully convolutional neural network by analyzing 14,503 T1 weighted structural MRI (sMRI) images of UK biobank data and achieved an MAE of 2.14 years in brain age estimation as a part of the 2019 predictive analysis challenge for brain age prediction. The authors of [15] constructed a novel convolutional neural network model by adding more filters in the initial phase of the architecture for more accurate brain age estimation (N=1445 T1 weighted sMRI images). These prior brain age studies mostly focused on predicting brain age using either single- or multi-modal structural or functional MRI data. However, the generalizability or robustness of these brain age models has not been comprehensively tested using independent samples.

One primary goal of this study is to test the replicability and generalizability of the pre-trained brain age models built in our previous study [1]. These brain age models were trained with the PNC dataset [20], consisting of 1417 adolescent subjects aged 8-22 years. Our previous study has shown that a traditional machine learning regression model with a small set of selected important features from a multimodal dataset can achieve high brain age prediction accuracy, similar to or better than those from complicated deep learning neural network models. In the current work, we investigated how previously trained brain age models perform on the Adolescent Brain and Cognitive Development (ABCD) [21–23] cohort. The ABCD participants have a narrow age range (9-10 years at baseline and 11-12 years at year-two follow-up). Furthermore, we also developed a refined brain age model that helps improve brain age models’ generalizability and prediction accuracy.

The rest of the paper is organized as follows. The materials and methods section describes the cohort, data preprocessing, and feature extraction procedure. The data analyses section presents detailed analytical approaches to evaluate PNC pre-trained brain age models, newly trained ABCD brain age models, and refined brain age models, followed by the association tests between cognitive ability and the brain age gap, and identification of the vital brain regions for brain age prediction. The test results and visualization of important brain regions are presented in the Results section. Finally, we provided the insights and knowledge we learned from our results in the Discussion and Conclusion section.

## 2 MATERIALS AND METHODS

### 2.1 Participants

The ABCD study (https://abcdstudy.org/) is a longitudinal study intended to run for 10 years. Over 11 thousand children at 9-10 years were recruited from 21 sites across the USA to collect a wide range of data, including brain imaging data, genetics, cognitive and behavioral assessments, family history and environmental measures, etc. The institutional review board approved protocols were followed to obtain all participating children’s consent and their parent’s full written informed consent. The current study used the available data from ABCD release 2.0, including baseline and year-two follow-up. Specifically, we have analyzed T1-weighted sMRI images of 11573 children from the ABCD baseline and T1-weighted sMRI images of 2947 children aged 11-12 years from the ABCD year-two follow-up.

### 2.2 Data preprocessing

We preprocessed sMRI images using the Statistical Parametric Mapping 12 (SPM12) software to generate the gray matter density maps. SPM12 jointly segments and spatially normalizes the T1 MRI data using SPM12 default tissue probability maps and generates six types of tissue maps (grey matter, white matter, CSF, bone, soft tissue, and others) in the Montreal Neuroimaging Institute (MNI) space. The gray matter maps were further smoothed by a 6 *mm*^3^ Gaussian kernel and resliced to a voxel size of 1.5 × 1.5×1.5 *mm*^3^. Quality control was applied to individual gray matter maps to select those correlated to the group mean gray matter map greater than 0.9, which resulted in 11,382 participants’ images in baseline (male: 5,960, female: 5,422; age: 9.92+/-0.63), 2877 in year-two follow-up (male:1553, female:1324; age: 11.93+/-0.63). Then we applied the same gray matter mask from our previous PNC study [1], on ABCD data to include the voxels with gray matter density larger than 0.2.

### 2.3 Feature extraction

The same sets of brain features used in our previous PNC brain age model were extracted for the ABCD cohort at baseline and year-two follow-up from preprocessed sMRI data. Specifically, we extracted three sets of brain structural features: 1) gray matter loadings of 100 brain components derived by independent component analysis (ICA) [24], 2) gray matter density of 116 regions of interest (ROI) based on the automatic anatomical labeling (AAL) [25] atlas, and 3) 152 anatomical features extracted by Freesurfer v5.3, software [26]. First, we have projected the same 100 ICA brain components used in our PNC brain age model onto ABCD baseline and two-year follow-up data. In our previous work, we applied the ICA model: *X*_*PNC*_ = *A*_*P NC*_ *×S*_*P NC*_. *X*_*PNC*_ denotes the gray matter data matrix, *A*_*PNC*_ is the loading matrix with each column indicating the expression of each component on individual subjects, and *S*_*P NC*_ is the 100 independent brain sources or component matrix of PNC cohort. Here we applied the projection equation 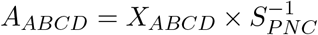 onto the ABCD cohort data to get the new loading matrix, *A*_*ABCD*_, which shows how the previous 100 ICA components derived from PNC data are expressed in the new ABCD cohort. It was conducted separately for ABCD baseline and year-two follow-up data. Therefore, the matrix, *A*_*ABCD*_, presenting loadings of gray matter components contains the feature vectors used for further brain age estimation. Secondly, we have used the AAL atlas map to extract the mean gray matter density features of 116 brain regions from the ABCD cohort. Thirdly, the ABCD study has provided tabulated data including FreeSurfer-derived imaging anatomical features for both baseline and year-two follow-up [23]. We selected the 152 anatomical features, which include estimated total intracranial volume, cortical thickness, and cortical and subcortical volumes of the left and right hemispheres of the brain based on the Desikan atlas.

## 3 Data Analyses for Brain Age models

### 3.1 PNC pre-trained brain age models

Our previous study was conducted using the data collected from PNC with adolescents aged 8-22 years. The primary goal of our PNC study was to utilize multimodal brain imaging data for brain age prediction and provide a detailed comparison of the brain age models’ performance trained on different unimodal or multimodal imaging modalities. Four types of imaging features were analyzed, including the three feature sets from gray matter images focusing on the brain structural features and one from the resting-state functional MRI – the loadings of 100 ICA components from derived functional network connectivity strength. To estimate the brain age, two sets of regression models, support vector regression (SVR, linear kernel) and partial least square regression with and without recursive feature elimination (RFE) were trained and tested using the brain imaging features as the independent predictors and the subject’s chronological age as the dependent variable. Details of the model building can be seen in reference [1]. In summary, we found that the SVR models with multimodal data and feature selection produced the best prediction accuracy. The multimodal SVR model trained with only 120 features selected from brain structural feature sets achieved an MAE of 1.25 years. The SVR model with selected 188 features from all four feature sets obtained an MAE of 1.22 years. Since the inclusion of the features from functional MRI data in the multimodal brain age model only provided limited improvement in the prediction accuracy, we focus on testing the reliability and generalizability of the brain age models only using structural brain features in the current study by excluding the functional data.

In this current research, we tested the performance of the pre-trained SVR brain age models with the ABCD baseline and year-two follow-up data. To comprehensively understand model reliability (e.g., whether individual feature sets have different reliability), we investigated eight brain age models listed in Table 1, named based on the features used. Table 1 summarizes the brain features used for individual models and their accuracy on PNC data reported in the previous study. For simplicity, we will refer to each brain age model as the short name in Table 1. The ABCD subjects have a narrow age range of 9-10 years and 11-12 years for baseline and year-two, respectively. We first evaluated all PNC pre-trained SVR brain age models using ABCD brain structural features at baseline, and then evaluated *FreeSurfer*_152, *FreeSurfer*_*RFE*_110, *Combined*_368, and *Combined*_*RFE*_120 models using ABCD year-two follow up data as these four models showed reasonable performance for baseline data as explained later. For the same reason, feature sets from these four models were used to build new models explained in the following as a comparison.

### 3.2 ABCD self-trained brain age models

To verify the predictive power of the ABCD data and to put the performance of pre-trained models into perspective, we newly trained four brain age models using FreeSurfer or combined brain features from ABCD baseline data. These four models used the same feature sets as in *FreeSurfer*_152, *FreeSurfer*_*RFE*_110, *Combined*_368, and *Combined*_*RFE*_120. We divided the baseline samples into 90% training and 10% testing data. Then on the 90% training data, we applied 10-fold cross-evaluation with the grid-search method to select the best hyper-parameters and best-fitted SVR models. The newly trained ABCD brain age models were tested on the 10% hold-out testing data, the full baseline samples, and the year-two follow-up samples. When directly testing the replication of PNC pre-trained models, we applied the models to the full samples of ABCD baseline and year-two follow-up. To make comparisons fair, we thus focused on the test performance of ABCD self-trained models and later refined brain age models using the full samples of ABCD baseline and year-two follow-up. We have reported the mean of predicted brain age, Pearson correlation coefficient (r-value) between actual age and predicted brain age, and MAE in years of predicted brain age to evaluate the performance of various brain age models.

### 3.3 Refined brain age models

We constructed refined brain age models (see Figure 1) to improve the robustness of existing brain age models. The refined model first leveraged the broader prediction power of PNC pre-trained models by feeding imaging features to the PNC pre-trained models. Then the refined model added a new module to capture the unexplained residuals, which are the differences between the actual age and the predicted age from the broader brain age models. Finally, the refined brain age was obtained by subtracting the estimated residuals from the broader predicted age. The model can be formulated as Equation 1 with the training of *g*(*X*). X is the input image features, the same set of features from the selected four models aforementioned. f(X) represents the PNC pre-trained model to compute 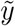, the estimated broad brain age. *g*(*X*) is a SVR module trained to explain the difference between 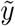 and chronological age, which presents the subtle brain age variation within a narrow age range. y is the final refined brain age.

**Figure 1.**
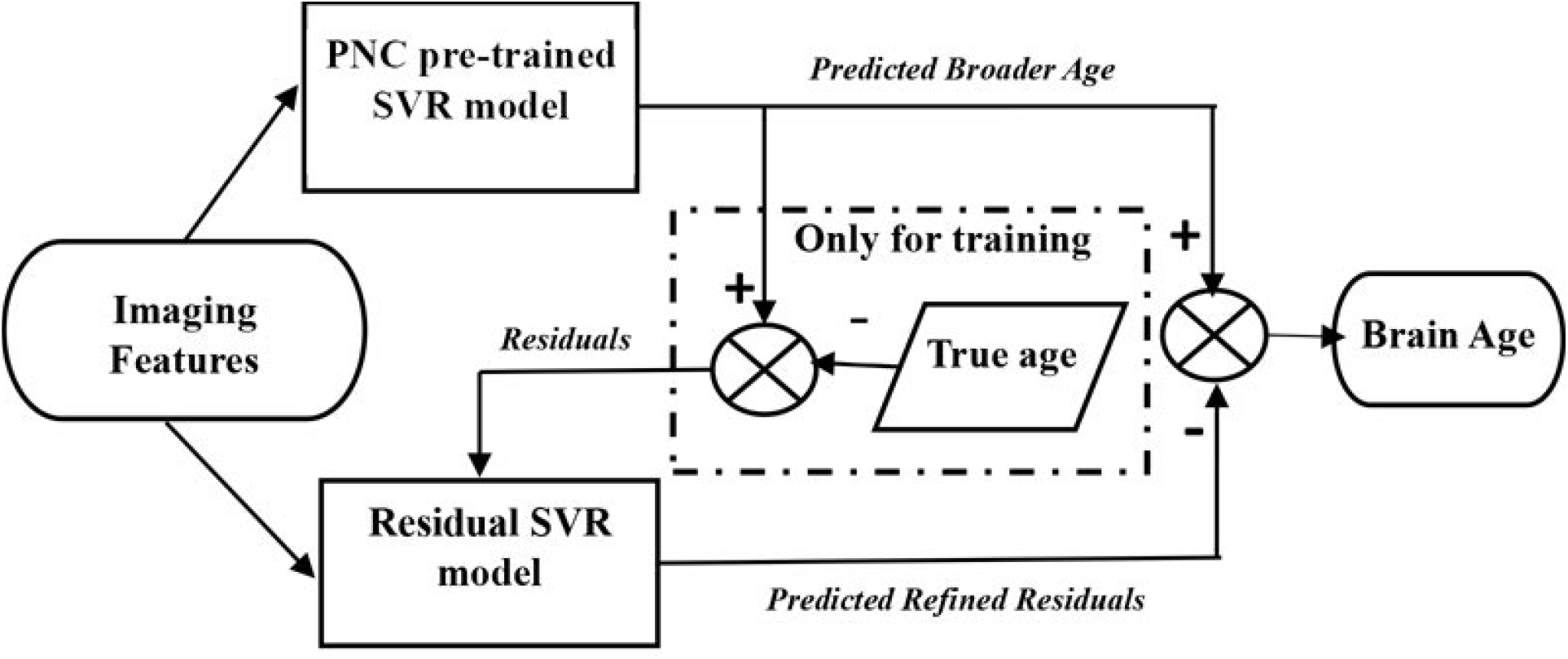
: Refined brain age model.

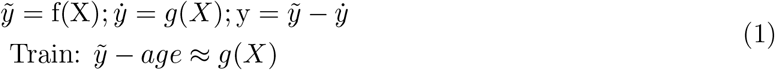

To train the refined models, we applied 10-fold cross-validation and grid-search hyper-parameter selection approach on 90% training data from ABCD baseline data. The performance of the refined brain age models was evaluated on 10% testing data of baseline, the full baseline data, and the full year-two follow-up data. Similarly, we only focused on the test performance on the full ABCD baseline, and year-two follow-up data to maintain the evaluation consistency among the brain age models.

### 3.4 Associations of brain age gap with cognitive measures

The brain age gap (the difference between the estimated brain age and the actual age) is calculated from the best-performing model. Then we examined the associations between individuals’ brain age gap and their cognitive ability. Six NIH toolbox cognitive measures were investigated separately. They are scores from (i) Flanker attention task to test the speed and accuracy of participants’ ability to identify congruent or incongruent targets given surrounding stimuli, (ii) pattern comparison processing speed task where the participants identify whether the pictures are the same or not from two given images and the score is calculated based on the number of correct answers, (iii) picture vocabulary task where the participants match the correct picture from the given four images by hearing an audio concept, (iv) picture sequence task where the participants replicate an activity sequence from a presented series of fifteen images that illustrate the particular activity, (v) the oral reading recognition task where the participants pronounce a single letter or word displayed in the middle of an iPad screen, and (vi) crystallized cognition composite score composed of picture vocabulary and oral reading recognition test.

These test scores indicate an individual’s executive function or attention ability, visual information processing speed, verbal intellect, episodic memory or visuospatial sequencing memory, language/academic achievement, and composite crystallized intelligence. We used age-uncorrected standard scores to reflect age-related cognitive development. Linear mixed effect models were implemented with cognitive measures as the dependent variable, sex (male=0, female=1), age, and brain age gap as fixed effect independent variables, and ABCD family ID nested with ABCD site information as random effect variables. We included each subject’s actual age and gender as predictors along with the brain age gap, as we want to control for the known age and gender effect on cognition during development. For investigating the proportion of variance uniquely explained in the response variable (cognitive measure) by the fixed effect predictor (brain age gap), we have calculated adjusted *R*^2^ following the approach of Stoffel et al. [27]. The adjusted *R*^2^ is calculated by the difference between the variance explained by the entire model, including all predictors, and the variance explained by a reduced model without the predictor of interest (brain age gap). We also reported a standardized coefficient value which is calculated by *β ×*(*sd*(*x*)*/sd*(*y*)), where *β* is the regression coefficient, x is the predictor and y is the response variable. Finally, we reported the p-values and corrected p-values (q-values) by multiple comparison corrections using the false discovery rate (FDR) method.

### 3.5 Important brain regions for age prediction

We have applied Shapley Additive Explanation (SHAP) technique [28] on our best-performing brain age model, refined *Combined*_*RFE*_120 model, to interpret the features’ contribution and to identify top brain areas which are crucial for brain age prediction. SHAP is a game theory-based method that computes the Shapley value for each feature by estimating the difference in the model outcome between a given sample and the averaged outcome across all data inputs. It reveals the individual feature contribution towards the model outcome for each sample or instance.

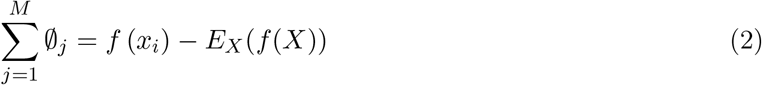

Here in the equation 2, *x*_*i*_ denotes one sample from a given dataset, *X* denotes the whole given dataset and ∅_*j*_ represents the SHAP value or feature importance of single feature *j*. The model outcome for a single sample is noted as *f* (*x*_*i*_) and *E*_*X*_(*f* (*X*)) denotes averaged outcome value or expected value on the whole sample. The difference between the model outcome of one sample and the model expected value equals the sum of all feature importance for the sample.

For linear models, with independent features, we can approximate SHAP values directly from the model’s weight coefficients. Given a linear model 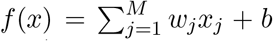 the importance or SHAP value can be calculated as : ∅ _j_ = w_j_ (x_j_ *−*E [X_j_]) where ∅_j_ is SHAP value, w_j_ and x_j_ represent regression coefficient and input value of feature *j* for the instance, respectively. E [X_j_] refers to the expected value for *X*_*j*_. We calculated the weights of the refined Combined_RFE_120 model using the following formula.

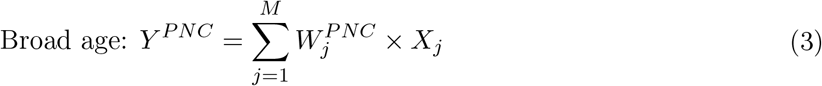

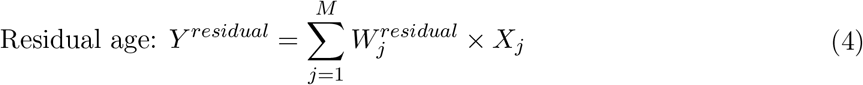

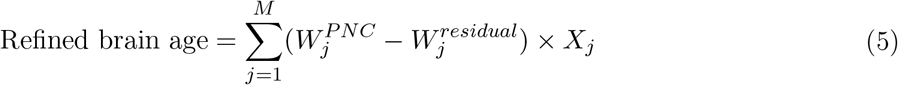

Here in Equation 3, *Y* ^*P NC*^, *W*^*P NC*^, and *X* indicate the output, model weights, and 120 RFE features of the first module (PNC pre-trained model) of our refined model, respectively. In Equation 4, *Y* ^*Residual*^ and *W*^*residual*^ correspond to the output and model weights of the second module (residual refinement) respectively. Finally, we calculated the refined model weights for each feature by subtracting the weights of the residual module from the weights of the PNC pre-trained module. We employed the linear SHAP method for the refined Combined_RFE_120 model and approximated the SHAP value of each feature. The 120 features were ordered based on the mean absolute value of SHAP values across all subjects. We have identified the top eight contributing features for brain age prediction by considering the threshold value *Z* [*mean*(|*SHAP*|)] *>* 1.96 (95% interval), where Z refers to the z-score of the mean absolute value of SHAP values. We then plotted the top eight important brain features for visual illustration purposes.

## 4 RESULTS

The performance of various brain age models is listed in Table 2 and Table 3 for ABCD baseline data and year-two follow-up data, respectively. The performance metrics include the mean of predicted brain age, correlation (r) with predicted brain age and actual age, and MAE of predicted brain age.

**Table 1:** Description of the Brain age models.

| Pre-trained SVR Models | Features | Accuracy on PNC data |  |
| --- | --- | --- | --- |
|  |  | *Train*<br>(MAE) | *Test*<br>(MAE) |
| ICA_100 | Loadings of 100 ICA components from gray matter images | 1.69 ± 0.23 | 1.68 |
| ICA_RFE_81 | RFE selected 81 ICA components | 1.64 ± 0.25 | 1.60 |
| ROI_116 | Gray matter density of 116 AAL brain regions | 1.77 ± 0.18 | 1.90 |
| ROI_RFE_70 | Gray matter density of RFE selected 70 AAL brain regions | 1.76 ± 0.19 | 1.73 |
| FreeSurfer_152 | 152 brain anatomical features from FreeSurfer | 1.77 ± 0.14 | 1.76 |
| FreeSurfer_RFE_110 | RFE selected 110 brain anatomical features from FreeSurfer | 1.70 ± 0.30 | 1.68 |
| Combined_368 | 368 combined brain structural features: 100 ICA components, gray matter density of 116 AAL regions, 152 brain anatomical features | 1.54 ± 0.22 | 1.32 |
| Combined_RFE_120 | RFE selected 120 combined brain structural features: 40 ICA components, mean density of 38 AAL brain regions, 42 anatomical features | 1.32 ± 0.13 | 1.25 |

**Table 2:** Evaluation of PNC pre-trained brain age models (Top), ABCD self-trained brain age models (Middle), and refined brain age models (bottom) using ABCD Baseline data.

| Brain Age Models | Model Name | Results |  |  |
| --- | --- | --- | --- | --- |
|  |  | Baseline (N= 11,382) |  |  |
|  |  | *Mean* | *Correlation (r)* | *MAE* |
| PNC Pre-trained | ICA_100 | 13.48 | 0.02 | 3.58 |
|  | ICA_RFE_81 | 12.75 | 0.03 | 2.89 |
|  | ROI_116 | 8.08 | 0.09 | 2.42 |
|  | ROI_RFE_70 | 7.70 | 0.08 | 2.66 |
|  | FreeSurfer_152 | 9.85 | 0.22 | 1.42 |
|  | FreeSurfer_RFE_110 | 9.76 | 0.22 | 1.43 |
|  | Combined_368 | 9.33 | 0.13 | 1.57 |
|  | Combined_RFE_120 | 9.53 | 0.13 | 1.56 |
| ABCD self-trained | FreeSurfer_152 | 9.91 | 0.34 | 0.50 |
|  | FreeSurfer_RFE_110 | 9.90 | 0.33 | 0.50 |
|  | Combined_368 | 9.91 | 0.45 | 0.46 |
|  | Combined_RFE_120 | 9.90 | 0.38 | 0.49 |
| Refined | FreeSurfer_152 | 9.89 | 0.17 | 0.76 |
|  | FreeSurfer_RFE_110 | 9.91 | 0.16 | 0.78 |
|  | Combined_368 | 9.91 | 0.21 | 0.60 |
|  | Combined_RFE_120 | 9.90 | 0.36 | 0.49 |

**Table 3:** Evaluation of PNC pre-Trained brain age models, ABCD self-trained brain age models, and Refined brain age Models using ABCD Year-Two Follow-Up data.

| Brain Age Models | Model Name | Results |  |  |
| --- | --- | --- | --- | --- |
|  |  | Two years follow-up (N= 2877) |  |  |
|  |  | *Mean* | *Correlation (r)* | *MAE* |
| PNC Pre-trained | FreeSurfer_152 | 11.11 | 0.26 | 1.56 |
|  | FreeSurfer_RFE_110 | 10.88 | 0.27 | 1.64 |
|  | Combined_368 | 10.38 | 0.16 | 1.92 |
|  | Combined_RFE_120 | 10.88 | 0.16 | 1.69 |
| ABCD self-trained | FreeSurfer_152 | 10.18 | 0.33 | 1.75 |
|  | FreeSurfer_RFE_110 | 10.18 | 0.33 | 1.75 |
|  | Combined_368 | 10.33 | 0.42 | 1.60 |
|  | Combined_RFE_120 | 10.22 | 0.37 | 1.72 |
| Refined | FreeSurfer_152 | 11.94 | 0.20 | 0.76 |
|  | FreeSurfer_RFE_110 | 11.93 | 0.21 | 0.74 |
|  | Combined_368 | 11.93 | 0.26 | 0.58 |
|  | Combined_RFE_120 | 11.92 | 0.40 | 0.48 |

Table 2 (top panel) from direct replication tests of PNC pre-trained models on ABCD baseline data showed that the Freesurfer and combined pre-trained models gave relatively more accurate brain age prediction than the ICA and ROI models in terms of predicted mean age. ICA and ROI models resulted in either overestimated or underestimated brain age estimation. The predicted age from all models showed correlations with actual age ranging from 0.02 to 0.22, and MAE between 1.42 to 3.58 years. Considering participants of ABCD at baseline were all 9-10 years old, the relatively good-performed models, including *FreeSurfer*_152, *FreeSurfer*_*RFE*_110, *Combined*_368, and *Combined*_*RFE*_120, were selected for further testing on year-two follow-up. They were also selected as bases for further model building for independently newly trained ABCD models and for refined brain age models. Table 2 (middle panel) lists the results of ABCD self-trained models. The four models presented similar performance with a predicted mean age of 9.9 years old, correlation from 0.34 to 0.45, and MAE of 0.46 to 0.50. The Combined_368 model was slightly better than others. Table 2 (bottom panel) displayed the performance of four refined brain age models on the full baseline samples, where the model with combined features showed relatively better performance. The *Combined*_*RFE*_120 model displayed the best performance with predicted age mean of 9.90, correlation of 0.36, and MAE of 0.49 years.

Table 3 presents the testing results on ABCD year-two follow-up data. Table 3 (top) showed the direct replication tests on four models with FreeSurfer and combined features, and all four models underestimated age at different levels, with mean age from 10.38 to 11.11. The correlation with actual age was from 0.16 to 0.27 and MAE from 1.56 to 1.92 years. Table 3 (middle) showed the performance of four ABCD self-trained models and all four models failed with various levels of underestimation of brain age. The predicted mean age was 10.38 to 11.11, and the MAE was 1.60 to 1.75. Table 3 (Bottom) listed the performance of refined models, and all four models achieved predicted mean age between 11.92 and 11.94 and MAE between 0.48 and 0.76. The best-performing model *Combined*_*RFE*_120 produced a mean of 11.92, a correlation of 0.40, and an MAE of 0.48.

The best-performing model across baseline and follow-up data is the *Combined*_*RFE*_120 refined model. Therefore, focusing on the *Combined*_*RFE*_120 refined model, we tested the relationship of the brain age gap with cognitive measures at baseline and year-two follow-up. Table 4 lists the standardized coefficient value, adjusted *R*^2^, p-values, and adjusted p-values (q-values) from association tests via linear mixed models. The significant results passing FDR correction are listed in bold. The positive coefficient value indicates that the subject’s cognitive ability increases with accelerated brain age.

**Table 4:**
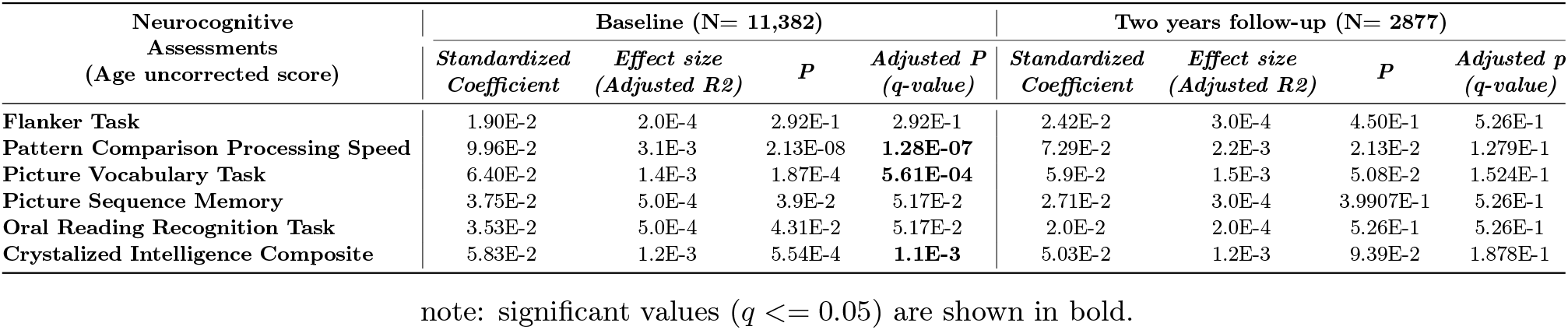
Associations of brain age gap from the refined *Combined*_*RFE*_120 model with cognitive measures at baseline and year-two follow-up.

We have also identified the top 8 contributing features of *Combined*_*RFE*_120 refined model using SHAP values as plotted in Figure 2. Figure 2 (a) shows a beeswarm plot indicating the top eight contributing features’ impact on brain age prediction. Each dot denotes the SHAP value of one feature for one individual subject, where the SHAP value is along the x-axis. and the blue-purple color indicates the input feature value from low to high. Figure 2 (b,c) shows the brain areas involved in the selected top contributing features in the coronal and sagittal view, color-coded as listed in Figure 2 (d).

**Figure 2.**
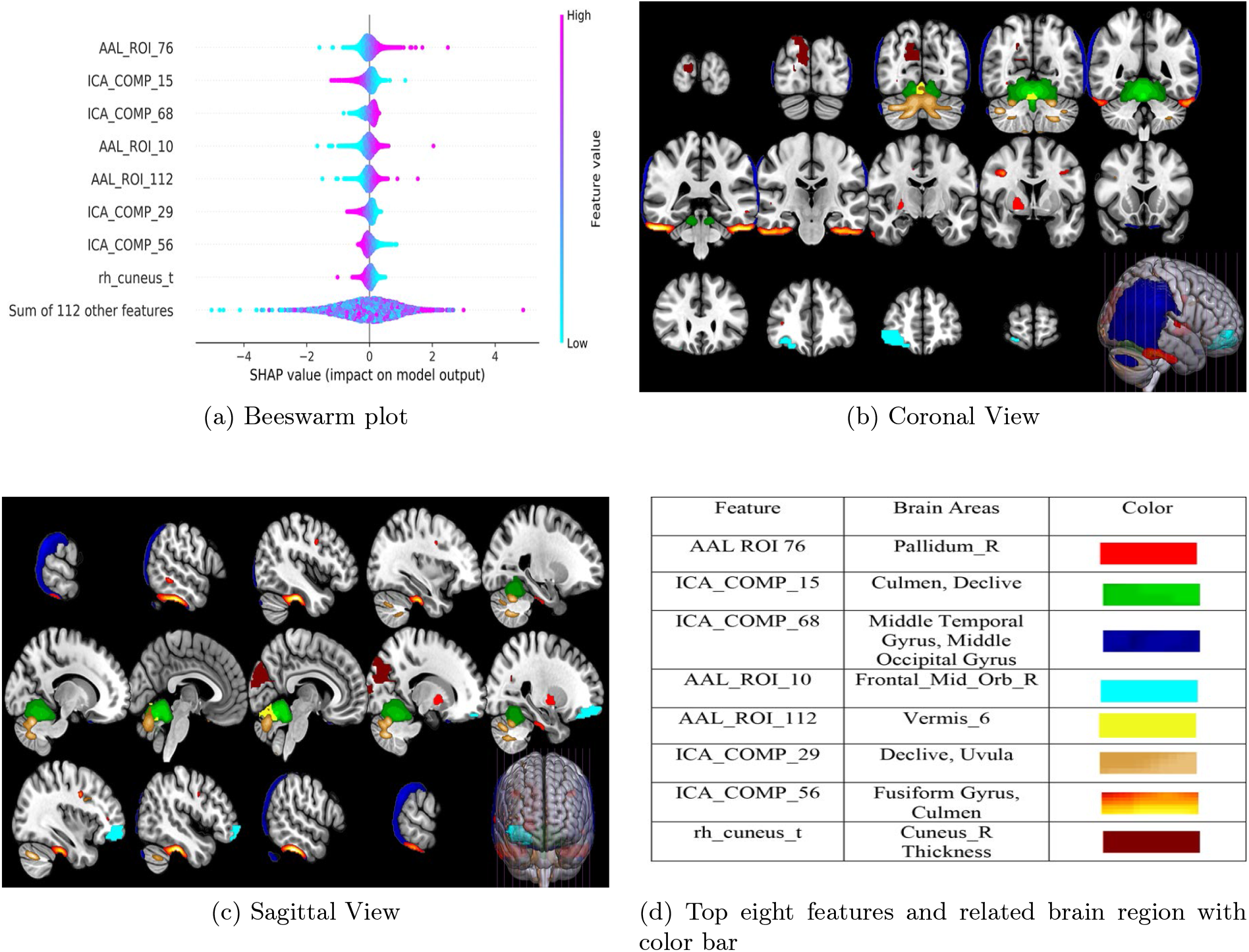
: Top eight features and the related brain areas from the refined model. a) The beeswarm plots showing the contribution (SHAP value) of the top eight features. b) Coronal view of top eight brain regions. c) Sagittal view of the top eight brain regions. d) Color code for the top eight features and their related brain locations.

## 5 Discussion and Conclusion

Recently many studies have used regression-based machine learning approaches or deep learning models to estimate brain age with high precision [29–32]. Very few studies focus on analyzing the generalizability of brain age prediction models. The study [32] shows that unseen data and different age ranges of samples can influence the generalizability of the brain age model. In this study, we evaluated the generalizability of the pre-trained brain age models (trained on PNC subjects with a broader age range of 8-22 years) from our previous study [1] by applying them to independent ABCD baseline and year-two follow-up data of children aged between 9-10 or 11-12 years. Our analysis showed that the model with only ICA component features or mean gray matter density features of brain regions showed lower accuracies in age prediction relative to the Freesurfer features. The models with ICA features produced overestimated predictions with low correlations with the actual age and high MAEs. In contrast, the models with mean gray matter density of ROIs underestimated the brain age with similar low correlations and high MAEs. We have followed the same preprocessing pipeline for both PNC and ABCD structural MRI images. However, the different group mean age of the subjects (approx. 16 years for PNC data, 9.9 and 11.93 years for ABCD baseline and year-two data, respectively) of PNC and ABCD cohorts might have induced some variability during the image spatial-normalization phase of preprocessing steps. That might be a possible reason behind the low performance of the ICA and AAL gray matter density ROI models. We have observed that the models with FreeSurfer and combined features achieved relatively better performance, with predicted brain age being closer to the mean of actual age from 9.33 to 9.85 at baseline and 10.38 to 11.11 at year-two follow-up. But the MAE of predicted age was still high, from 1.42 to 1.57 at baseline and 1.56 to 1.92 at year-two, and the predicted age did not correlate strongly with the actual age on both baseline and year-two follow-up. Focusing on Freesurfer and combined features, we trained the new ABCD brain age models using the ABCD baseline data to test the prediction power of the ABCD features. As expected, the ABCD self-trained brain age models showed improved accuracy when tested on baseline data, but the performance decreased when tested on the year-two follow-up data. Basically, all four ABCD self-trained brain age models failed while tested on year-two follow-up data when participants were 11-12 years old. For instance, the performance of the *Combined*_368 model declined markedly with the MAE increasing from 0.46 at baseline to 1.92 at year-two follow-up.

To improve the generalizability of the brain age model, we have constructed refined brain age prediction approach. Our results indicate that the refined brain age models showed better prediction accuracy compared to both pre-trained models and self-trained models. In particular, for the *Combined*_*RFE*_120 refined model, the correlation with actual age significantly improved from 0.13 to 0.36, and the MAE decreased from 1.56 to 0.49 years for the baseline data (Table II). Similarly, we also observed big improvements for the year-two follow-up data for the *Combined*_*RFE*_120 refined model, when the correlation with actual age increased from 0.16 to 0.40 and the MAE decreased from 1.69 to 0.48 years (see Table 3). Unlike ABCD self-trained models, refined brain age models achieved improved performance on both baseline and year-two data.

Overall, the PNC pre-trained models failed to produce valid age prediction for both ABCD baseline and year-two follow-up, the ABCD self-trained models performed very well at baseline but failed at year-two follow-up, while the ABCD refined brain age models showed consistently good performance at both baseline and year-two follow-up. Even though all three types of models are linear, the weights are trained with different loss functions: minimize errors in a large age range, minimize errors in a narrow age range, and minimize errors in an adjusted narrow age range. The adjusted ages prevented the model to be over-fitted to the narrow age range.

For interpreting the meaning of the brain age gap, we have applied linear mixed-effect models to test the associations between the derived brain age gap and six neurocognitive measures. As expected, age had a significant impact on all six cognitive measures for both time points, and sex had a significant influence on episodic memory and information processing speed at baseline and year-two follow-up. Detailed results are not presented here since age and sex effect on cognition is outside the scope of this study. Beyond age and sex, there is a significant association between the brain age gap and the rate of information processing speed (pattern comparison), verbal comprehension (picture vocabulary), and crystallized intelligence on baseline data. The adjusted *R*^2^ score, partial variance explained is 0.0031, 0.0014, and 0.0012, respectively, indicating accelerated brain age explains partially, small yet significant, the advance in cognitive ability. For the year-two analysis, the uncorrected p-values were less than 0.05 for pattern comparison and 0.05 for the picture vocabulary task. Even though the association tests produced similar effect sizes with consistent positive effects between baseline and year-two follow-up, we did not observe any significant influence (based on FDR corrected q-value) of the brain age gap over the cognitive measures for the year-two data. It may be due to a comparatively smaller sample size of data (baseline: N=11,382, year-two: N=2877) in the year-two study.

Eight brain features are selected to be important for brain age prediction. Based on the features’ SHAP values (Figure 2(a)), we can interpret how the top features influence brain age. For example, the highest contributing feature, *AAL*_*ROI*_76, has SHAP values ranging from about -3 to 3. The positive SHAP values are presented in purple color, indicating higher input feature values, while negative SHAP values are in blue indicating lower input features. Thus, a higher value of the feature *AAL*_*ROI*_76 relates to an increased brain age. Vice versa, a lower feature value in *AAL*_*ROI*_76 links to a decreased brain age. *AAL*_*ROI*_76 is the gray matter volume of the right Pallidum shown in Figure 2(b,c). Therefore, our model suggests an increased gray matter density in the right pallidum area is a biomarker for the subject’s brain development/maturation during this sensitive developing phase. Similarly, the relationship between the SHAP value and the input value of the second top feature (*ICA*_*COMP* _15 positive SHAP values link to low input values) suggests lower gray matter density in the culmen area is associated with increased brain age. In sum, our results suggest that the higher gray matter density in the right pallidum, right middle orbital frontal gyrus, middle temporal gyrus, and vermis is associated with increased brain age, while the lower gray matter density in the culmen, declive and fusiform gyrus, as well as reduced cortical thickness of Cuneus, is associated with increased brain age. It is well documented that gray matter develops in a region-specific manner with a ‘posterior to anterior’ maturation order at large [33], and presents both linear and non-linear (e.g., inverted U shape) developmental trajectories [34–36]. For instance, Wierenga and colleagues [37] focused on subcortical regions’ developmental trajectory and reported that pallidum presents a curvilinear trajectory peaked at 17 years old. Thus, at the age of 9 to 12, the increase of pallidum volume in the participants of the ABCD cohort indicates continued brain development and increased brain age. For the cortical area of the brain abundant evidence supports that higher-order association cortices mature only after lower-order somatosensory and visual cortices, with gray matter thickness peaking at different ages [33, 38]. At the age of 9-12, our model indicates brain development is linked to reduced gray matter density and thickness in the occipital lobe (fusiform gyrus, cuneus) and increased gray matter density in the frontal and temporal lobe, suggesting the visual cortex has passed its maturation peak but the higher order frontal and temporal cortices are still approaching to their peaks. In contrast, few studies have investigated cerebellum and cerebellar subregions development during childhood or life span [39, 40]. Consistently both studies identified that most of the cerebellar subregions followed an inverted U-shaped developmental trajectory and Romero et al. also highlighted the Lobule IX (the uvula) showed the fastest development, reaching the maximum at around 5 years [40]. Our model also makes a note of gray matter density in Uvula: reduced gay matter in Uvula indicates increased brain age at the age of 9-12. In addition, our model also grouped culmen, declive, and uvula, to have similar development change at this period of age. Overall, our findings indicate that in the pre-teens age, while cerebellum and occipital regions start gray matter reduction, the frontal and temporal regions are still in the increasing phase, which is in line with known brain development trajectory in children and adolescents [41, 42].

In summary, our study shows that the PNC pre-trained models have prediction power to capture the overall mean age of ABCD samples but cannot precisely estimate the brain age within the narrow range. In contrast, ABCD self-rained brain age models can only give a precise estimation of the subjects between 9-10 years but cannot be generalized to other age ranges. Our proposed refined age methods utilize the broader prediction power of pre-trained models and refined the model with residual to increase the overall robustness of the brain age models. The strength of this study should be interpreted in light of the following limitations. The two datasets tested (ABCD baseline and year-two follow-up) have an age range of pre-teens, and the robustness of the refined model requires more tests using data from teenagers and young adulthood, as well as data from independent samples other than ABCD. As the brain develops in a nonlinear manner during childhood and adolescence, the effectiveness of refined models may be subjected to periods of development, such as childhood, pre-teens, teenagers, and young adulthood. Future evaluation is warranted.

## Acknowledgment

This work is supported by NIH R01DA049238 and NSF 2112455 to VC and JL.

## Notes

### Competing Interest Statement

The authors have declared no competing interest.

